# The SARS-CoV-2 Spike Protein Mutation Explorer: Using an Interactive Application to Improve the Public Understanding of SARS-CoV-2 Variants of Concern

**DOI:** 10.1101/2022.09.09.507349

**Authors:** Sarah Iannucci, William T. Harvey, Joseph Hughes, David L. Robertson, Matthieu Poyade, Edward Hutchinson

## Abstract

SARS-CoV-2 is the virus responsible for the COVID-19 pandemic, which began in late 2019 and has resulted in millions of death globally. The need to understand the pandemic means that detailed descriptions of features of this virus are now of interest to non-expert audiences. In particular, there has been much public interest in the spike protein that protrudes from the surface of the SARS-CoV-2 virus particle. The spike is the major determinant of viral infectivity and the main target for protective immune responses, and included in vaccines, and so its properties influence the impact of the pandemic on people’s lives. This protein is rapidly evolving, with mutations that enhance transmissibility or weaken vaccine protection creating new variants of concern (VOCs) and associated sub-lineages. The spread of SARS-CoV-2 VOCs has been tracked by groups such as the COVID-19 Genomics UK consortium (COG-UK). Their online mutation explorer (COG-UK/ME), which analyses and shares SARS-CoV-2 sequence data, contains information about VOCs that is designed primarily for an expert audience but is potentially of general interest during a pandemic. We wished to make this detailed information about SARS-CoV-2 VOCs more widely accessible. Previously work has shown that visualisations and interactivity can facilitate active learning and boost engagement with molecular biology topics, while animations of these topics can boost understanding on protein structure, function, and dynamics. We therefore set out to develop an educational graphical resource, the SARS-CoV-2 Spike Protein Mutation Explorer (SSPME), which contains interactive 3D molecular models and animations explaining SARS-CoV-2 spike protein variants and VOCs. We performed user-testing of the original COG-UK/ME website and of the SSPME, using a within-groups design to measure knowledge acquisition and a between-groups design to contrast the effectiveness and usability. Statistical analysis demonstrated that, when compared to the COG-UK/ME, the SSPME had higher usability and significantly improved participant knowledge confidence and knowledge acquisition. The SSPME therefore provides an example of how 3D interactive visualisations can be used for effective science communication and education on complex biomedical topics, as well as being a resource to improve the public understanding of SARS-CoV-2 VOCs.

## 1 Introduction

Severe Acute Respiratory Syndrome Coronavirus 2 (SARS-CoV-2) emerged in late 2019, leading to the global Coronavirus Disease 2019 (COVID-19) pandemic which has claimed over 6.4 million lives as of August 2022 (World Health Organization, 2022). SARS-CoV-2 is a betacoronavirus with a ribonucleic acid (RNA) genome; of the proteins it encodes, four are incorporated into virus particles, namely the spike (S), envelope (E), membrane (M), and nucleocapsid (N) proteins (Peacock, et al., 2021; Chan, et al., 2020). The spike proteins are distributed across the outer viral envelope, giving the virus the characteristic crown-like or ‘solar corona’ appearance from which it gets its name (Ke, et al., 2020; Benton, et al., 2020). The spike protein is a homotrimer, meaning that it is composed of three identical smaller units called monomers; each monomer is split into two subunits, termed S1 and S2. The S1 subunit contains two key regions termed the N-terminal domain (NTD) and receptor-binding domain (RBD). As indicated by its name, the RBD binds to the host cell surface receptor Angiotensin-Converting Enzyme 2 (ACE-2) (Peacock, et al., 2021; Letko, et al., 2020) and is also the part of spike principally target by antibodies of the immune system. The N-terminal domain is also an important target for the immune system, to which host antibodies can bind to block infection (Casalino, et al., 2020). The spike protein can alternate between ‘open’ and ‘closed’ conformations, allowing for the RBD to bind to host cells when open, and avoid recognition by host antibodies when closed (Ke, et al., 2020; Benton, et al., 2020). The S2 subunit plays an important role in membrane fusion, where the virus fuses with the host cell to release its contents (Wrobel, et al., 2020). The spike proteins are one of the key determinants of infectivity and, because their functions can be blocked by the binding of antibodies, of antiviral immunity (Peacock, et al., 2021).

SARS-CoV-2 has undergone significant evolution since becoming established as a new human pathogen, with mutations that cause changes in the spike protein having particularly important consequences for the course of the pandemic. As the spike protein acquires changes in regions that improve the ability of the virus to infect, transmit, and evade the host’s immune system, such as the RBD and NTD, variants which are more harmful to the human population – termed variants of concern (VOCs) – have emerged (Harvey, et al., 2021). As of August 2022, five key variants or VOCs have been designated, referred to as the Alpha, Beta, Gamma, Delta, and Omicron (World Health Organization, 2022), of which Omicron is currently circulating around the globe. Many research groups and public health bodies are monitoring the spread of SARS-CoV-2 and its accumulating mutations, including the COVID-19 Genomics UK consortium (COG-UK). COG-UK maintains an online Mutation Explorer (ME) resource (COG-UK/ME, http://sars2.cvr.gla.ac.uk/cog-uk/), which provides open access data on SARS-CoV-2 spike mutations and VOCs (Wright, et al., 2021). As this resource is tailored to the needs of expert users, we recognised that its data are likely to be less accessible to a non-expert audience. We reasoned that incorporating interactive visual resources into the COG-UK/ME would allow us to combat this issue and enhance public understanding of the vital data contained on the website.

It has been demonstrated that introducing interactivity and visualisations can bring molecular biology topics into a more comprehensible format for those without a scientific background (Jenkinson, 2018). Interactivity can better allow the user to engage with the material, boosting ‘active learning’ and allowing for higher engagement and deeper information retention (Bruce-Low, et al., 2013). Including three-dimensionality into teaching molecular biology can also be informative, as it better allows the user to view protein structures and gain spatial cues, as well as boosting understanding of protein function (Höffler, 2010). Additionally, by visualising these complex topics in animations rather than in static formats, it is possible to convey protein dynamics with a visual narrative, which can further facilitate learning (Barak & Hussein-Farraj, 2013; Jenkinson, 2018). Therefore, user learning may be improved by incorporating interactive visual resources into resources conveying complex scientific information, such as the COG-UK/ME.

To address this need for interactivity, we created a graphical resource, the SARS-CoV-2 Spike Protein Mutation Explorer (SSPME). The SSPME is an online resource that brings together 3D models and narrated molecular animations into an interactive application to explain SARS-CoV-2 spike protein amino acid substitutions (linked to underlying point mutations in the virus RNA) and VOCs to a general audience. User-testing showed that SSPME provides a better user experience and increases the acquired knowledge of non-specialists when compared to the COG-UK/ME website. As well as providing a resource to help the general public interpret information about SARS-CoV-2 VOCs and to help the explanation of these topics in the media, this research provides a case study of using interactive graphical resources to facilitate molecular biology education for the general public.

## 2 Materials and Methods

### 2.1 The SARS-CoV-2 Spike Protein Mutation Explorer

Early user-testing of the COG-UK/ME was completed in order to compile feedback on this resource and identify areas for improvement that could be addressed within the SSPME. Participants from this early user-testing of the COG-UK/ME would form the Control Group (CG) in this research. CG results indicated that this development should include more introductory information on the SARS-CoV-2 viral structure, including more information about the regions of the spike protein that were most relevant for VOCs (see Supplementary Material). In addition, engaging visualisations were requested for use throughout the COG-UK/ME website in order to facilitate user understanding of these concepts.

A combination of visualisation techniques were used to develop the SSPME, containing information on SARS-CoV-2 spike mutations and VOCs. Briefly, using a combination of protein visualisation and 3D modelling software (Table 1), 3D models of relevant proteins were created from experimentally-resolved protein structures made available through the RCSB Protein Databank (Table 2). An accurate, interactive spike protein model was developed for use in the application and within the animations. In addition, 3D models of other proteins relevant for SARS-CoV-2 properties (see Table 2) were created using mMaya and used within the animations. All scientific content developed within the animations and the application was reviewed and validated by virologists from the MRC-University of Glasgow Centre for Virus Research and COG-UK/ME. The final application, the SSPME, is hosted freely online (https://sc2-application.itch.io/sars-cov-2-mutation-explorer). The full methodology used to create the SSPME have been described in detail previously (Iannucci, et al., InPress).

**Table 1.** The protein visualisation and 3D modeling software used in the development of the SSPME.

| Software | Use | Developer |
| --- | --- | --- |
| Autodesk Maya | 3D modeling and animation | Autodesk, Inc.<br>(https://www.autodesk.com/) |
| Molecular Maya<br>(mMaya) | Importing, modeling, rigging,<br>and animating molecular<br>structures within Autodesk<br>Maya | Digizyme, Inc.<br>(https://www.digizyme.com/) |
| PyMOL | Molecular visualisation and<br>3D model generation | Shrödinger, Inc.<br>(https://pymol.org/) |
| UCSF Chimera | Molecular visualisation and<br>3D model generation | Resource for Biocomputing,<br>Visualization, and Informatics (RBVI), |
|  |  | UCSF<br>(https://www.cgl.ucsf.edu/chimera/) |

**Table 2.** The experimentally resolved data used in generating the 3D models of proteins during the development of the SSPME.

| Protein | PDB/File Identifier | Reference |
| --- | --- | --- |
| Angiotensin-Converting Enzyme 2 (ACE-2) | 6M17 chains B, E | (Yan, et al., 2020) |
| Full-length glycosylated spike protein, 1 RBD open | 6VSB_1_1_1 | (Woo, et al., 2020) |
| Full-length glycosylated spike protein, all RBDs closed | 6VXX_1_1_1 | (Woo, et al., 2020) |
| Membrane protein | M_dimer_new_1 | (Heo & Feig, 2020) |
| Envelope protein | E_protein_1 | (Heo & Feig, 2020) |
| Lysosome-Associated Membrane Protein 1 (LAMP-1) | 5GV0 | (Terasawa, et al., 2016) |
| V-ATPase | 5VOX | (Zhao, et al., 2017) |
| Transient Receptor Potential Mucolipin 1 (TRPML1) | 5WJ5 | (Schmiege, et al., 2017) |
| Solution structure of human secretory IgA1 | 3CHN | (Bonner, et al., 2009) |
| Crystal Structure of Pembrolizumab, a full length IgG4 antibody | 5DK3 | (Scapin & Strickland, 2015) |

**Table 3.** The demographics collected from all participants within this study.

|  | Control Group | Test Group |
| --- | --- | --- |
| **Female participants** | 5 | 6 |
| **Male participants** | 4 | 8 |
| **Age range (Mean ± SD)** | 21-64 (42.78 ± 12.88) | 20-66 (41.14 ± 14.88) |
| **Level of attained education** | Range from secondary school (14%) to a Master's degree (57%) | Range from secondary school (7%) to a Ph.D. or higher (50%) |
| **Computer use** | Use a computer every day or almost every day | Use a computer every day or almost every day |
| **Self-reported experience with interactive applications** | 67% experienced | 64% experienced |
|  | 22% with some experience | 7% with some experience |
|  | 11% with no experience | 29% with no experience |
| **Self-reported general virology knowledge** | 14% knowledgeable | 65% expert or knowledgeable |
|  | 57% with a basic knowledge | 21% with a basic knowledge |
|  | 29% with no knowledge | 14% with no knowledge |
| **Self-reported SARS-CoV-2 knowledge** | 29% knowledgeable | 64% expert or knowledgeable |
|  | 57% with a basic knowledge | 36% with a basic knowledge |
|  | 14% with no knowledge | 0% with no knowledge |
| **Self-reported understanding of SARS-CoV-2 VOCs** | 29% well-informed | 50% well-informed |
|  | 28% moderately informed | 21% moderately informed |
|  | 43% slightly informed | 29% slightly informed |

The SSPME has the goal of providing the public with a concise and comprehensible visual resource for the SARS-CoV-2 spike protein and VOCs, as well as containing animations that explain key viral processes related to SARS-CoV-2 infectivity, transmissibility, and antigenicity. The features of the application are explained briefly below.

When a user visits the SSPME website, after encountering the opening and instruction pages, they are introduced to a 3D SARS-CoV-2 spike protein model embedded in a portion of the viral membrane. Using the computer mouse, the user can rotate, zoom in, and interact with four main regions of interest (ROI) on the spike protein: the RBD, the NTD, the FCS, and the glycans (Figure 1). From this area of the SSPME, the user can also alternate the spike protein’s conformation between open and closed (two different states that are important for understanding SARS-CoV-2 infection and immunity (Harvey, et al., 2021)), download a screenshot of the protein in its current position, access the VOC menu, and review a glossary panel of technical terms (Figure 2). The VOC menu provides an overview of VOCs that have played a dominant role in the pandemic up to the time of release (when the application was first released on 09-July-2021, this was the Alpha, Beta, Gamma, and Delta VOCs; the application was updated on 01-Dec-2021 to include the Omicron VOC) (World Health Organization, 2022). To illustrate the specific amino acid substitutions that define these VOCs clearly, we used static 3D visualisations provided by the COG-UK/ME (Wright, et al., 2021).

**Figure 1.**
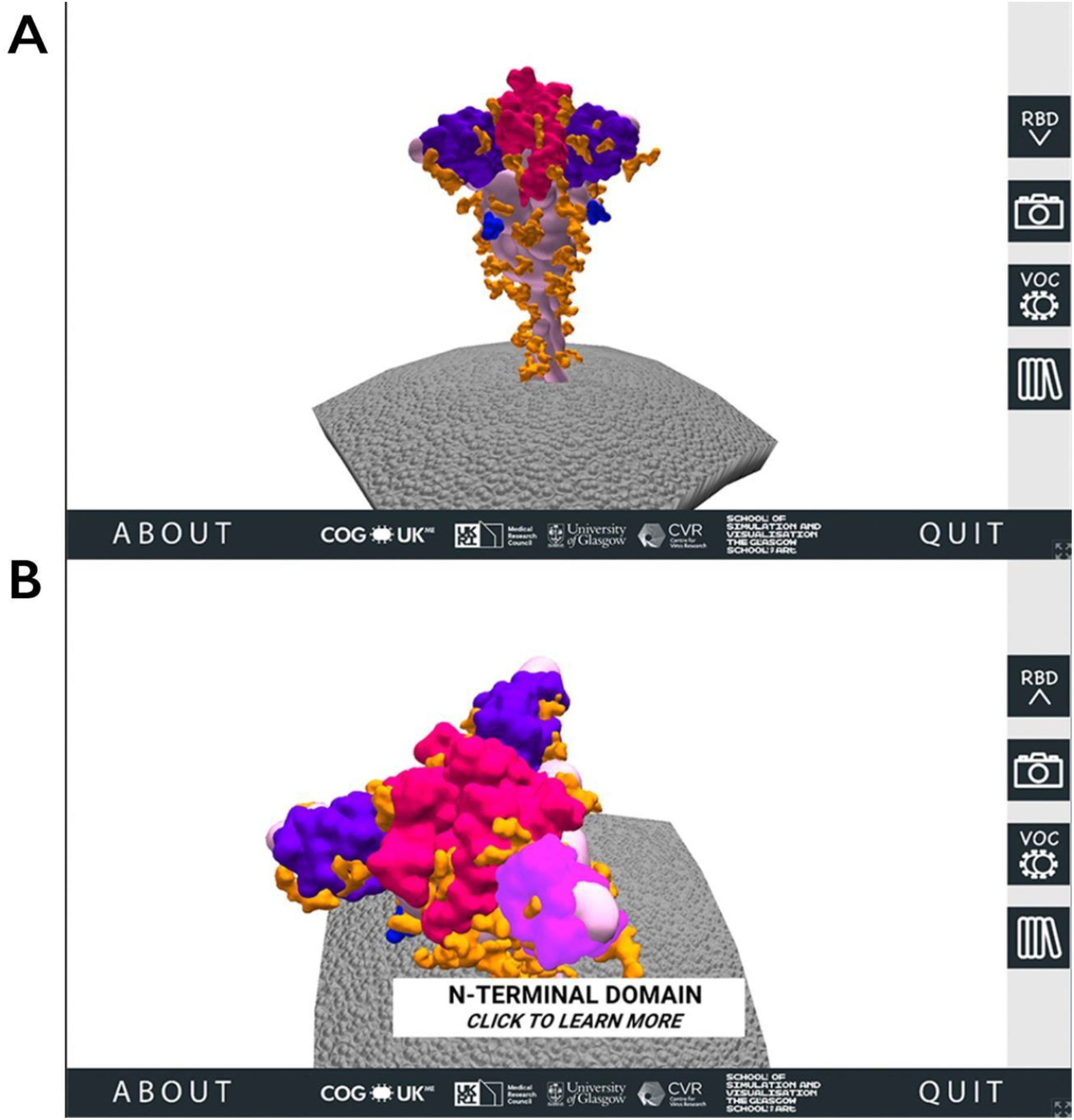
The 3D SARS-CoV-2 spike protein scene of the SARS-CoV-2 Spike Protein Mutation Explorer. Screenshots show the 3D SARS-CoV-2 spike protein embedded in the viral membrane (A) and the same scene when the user has rotated the model and is hovering their mouse over the NTD (B).

**Figure 2.**
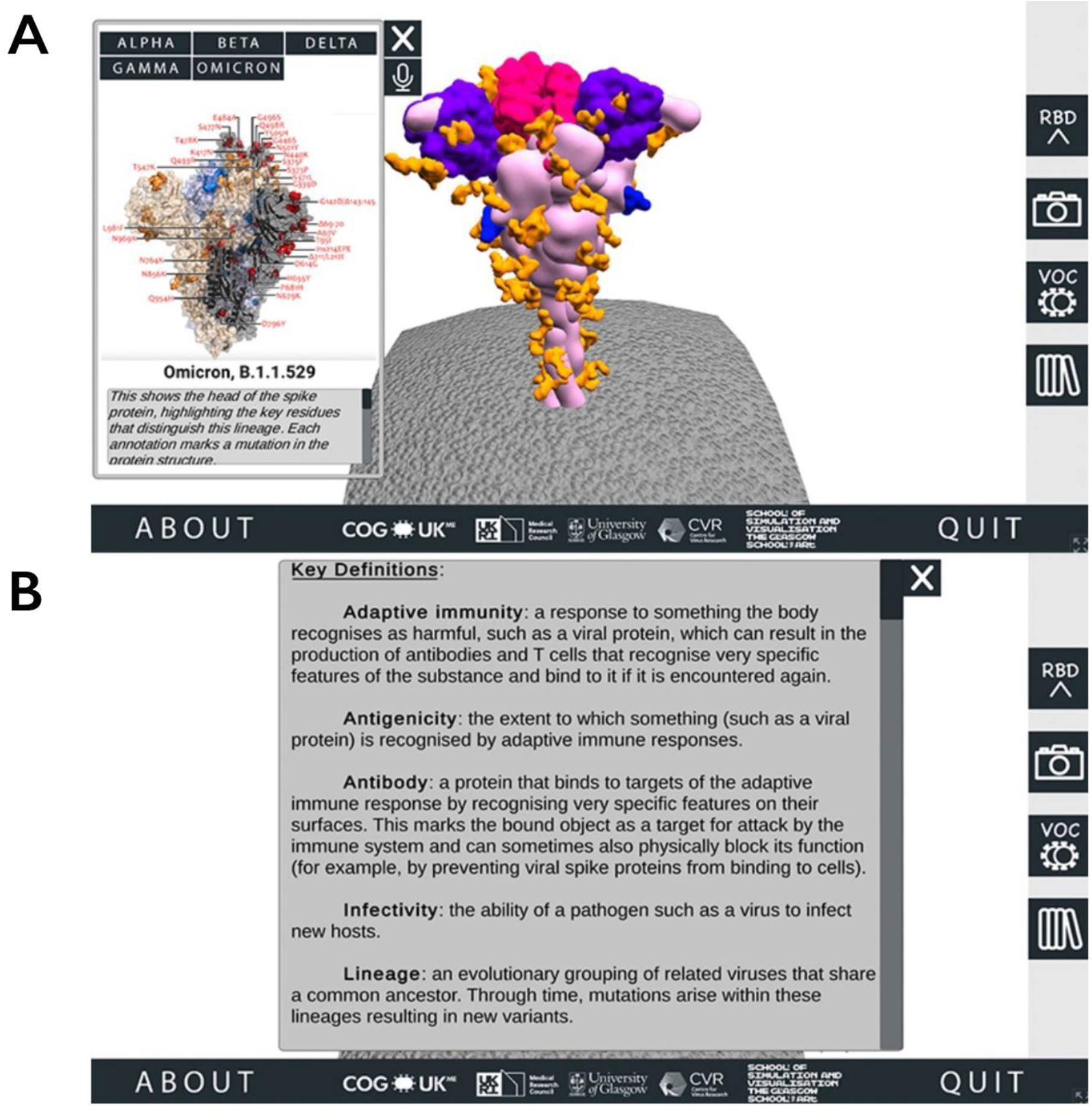
Accessing additional information through the SARS-CoV-2 Spike Protein Mutation Explorer. Screenshots show (A) the VOC menu and (B) the Glossary panel.

To understand why these amino acid substitutions affect SARS-CoV-2 biology, users need to be provided with detailed contextual information about the ROIs in which they occur. When a user mouse-clicks a ROI on the spike protein, they are taken to that ROI’s corresponding region-specific page, which goes into more detail about the ROI and how changes in this region may impact the properties of SARS-CoV-2 (Figure 3).

**Figure 3.**
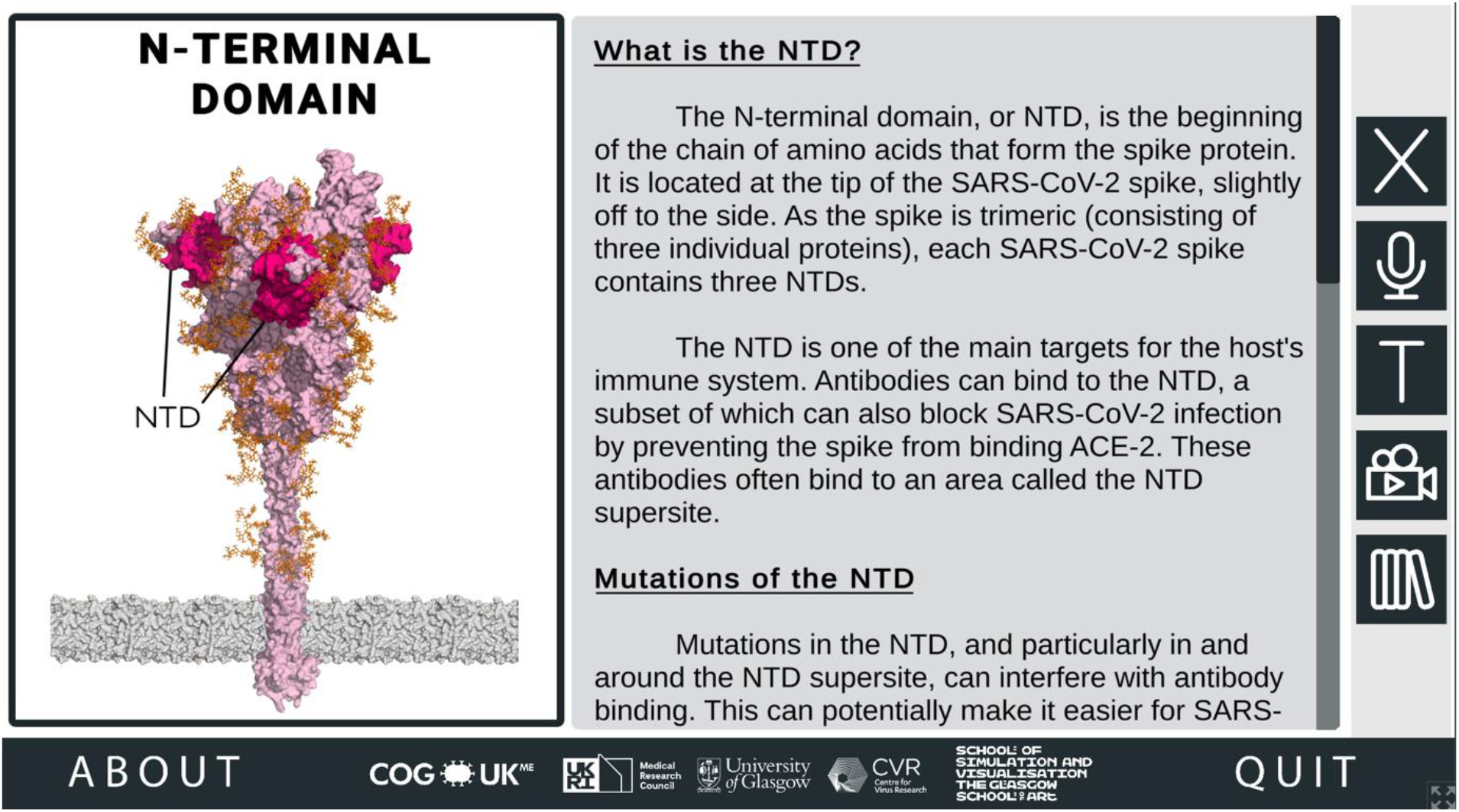
Information about Regions of Interest (ROI) in the SARS-CoV-2 Spike Protein Mutation Explorer. Screenshot shows the NTD-specific ROI page, one of the four ROI-specific pages.

It is from the RBD- and NTD-specific pages that the 3D molecular animations can be accessed, demonstrating key viral properties and providing visual explanations of how changes in these regions could affect SARS-CoV-2 (Figure 4).

**Figure 4.**
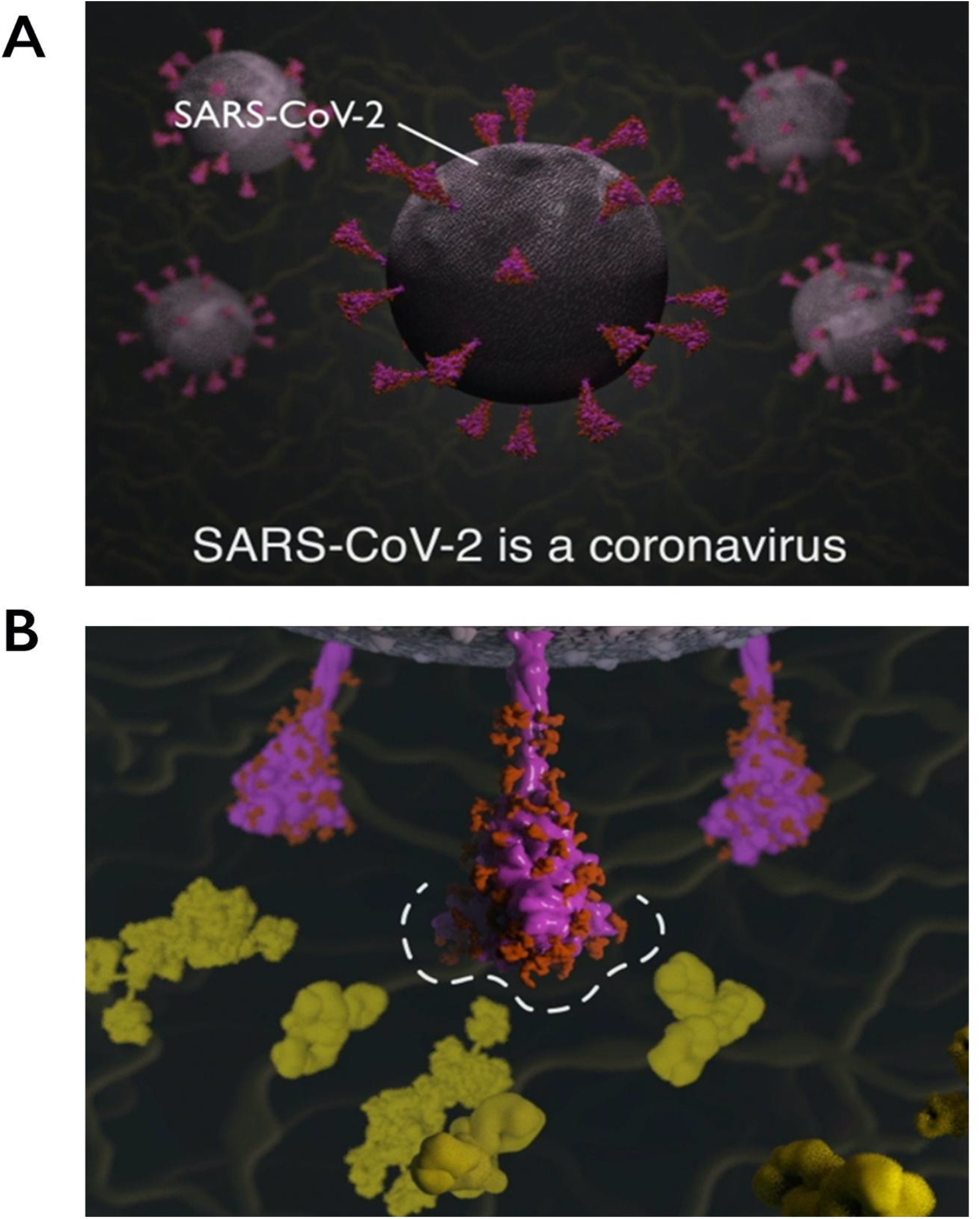
Explanatory animations in the SARS-CoV-2 Spike Protein Mutation Explorer. Screenshots show the 3D molecular animations contained within the RBD- and NTD-specific scenes of the SARS-CoV-2 Spike Protein Mutation Explorer, showing (A) SARS-CoV-2 virus particles and (B) a close-up SARS-CoV-2 spike protein avoiding host antibodies, with the antibody binding sites highlighted with a dashed line. Mucin proteins present in the nasal mucus are visible as background detail.

### 2.2 Experimental Methods

User-testing was carried out to compare the impact of the SSPME to that of the original COG-UK/ME site on learning and knowledge acquisition of the users, as well as exploring the usability of the application. This study received Ethical Approval from The Glasgow School of Art’s Learning and Teaching Office.

#### 2.2.1 Participant Recruitment and Demographics

In order to recruit participants for this study, digital posters were regularly advertised on Twitter, making particular use of the MRC-University of Glasgow Centre for Virus Research (https://twitter.com/CVRinfo) account which, due to their subject matter and high follower count, could potentially extend our reach for recruitment. Participants signed up via a Google Form which confirmed that they were 16 years or older and had access to a computer and internet. Participants were anonymised throughout the study and assigned to two groups: a Control Group (CG) which would examine the original COG-UK/ME website, and a Test Group (TG) which would examine the SSPME. Informed consent and demographics were collected from all participants (Table 1). All participant demographics were later used in the assessment of the user-testing phase of this study.

#### 2.2.2 Experimental Procedure and Data Analysis

Following participant recruitment, sign-up and obtainment of informed consent, participants were privately emailed an overview of the experimental study and given access to the experimental questionnaire (taking them through seven stages; see Supplementary Material) (Figure 5).

**Figure 5.**
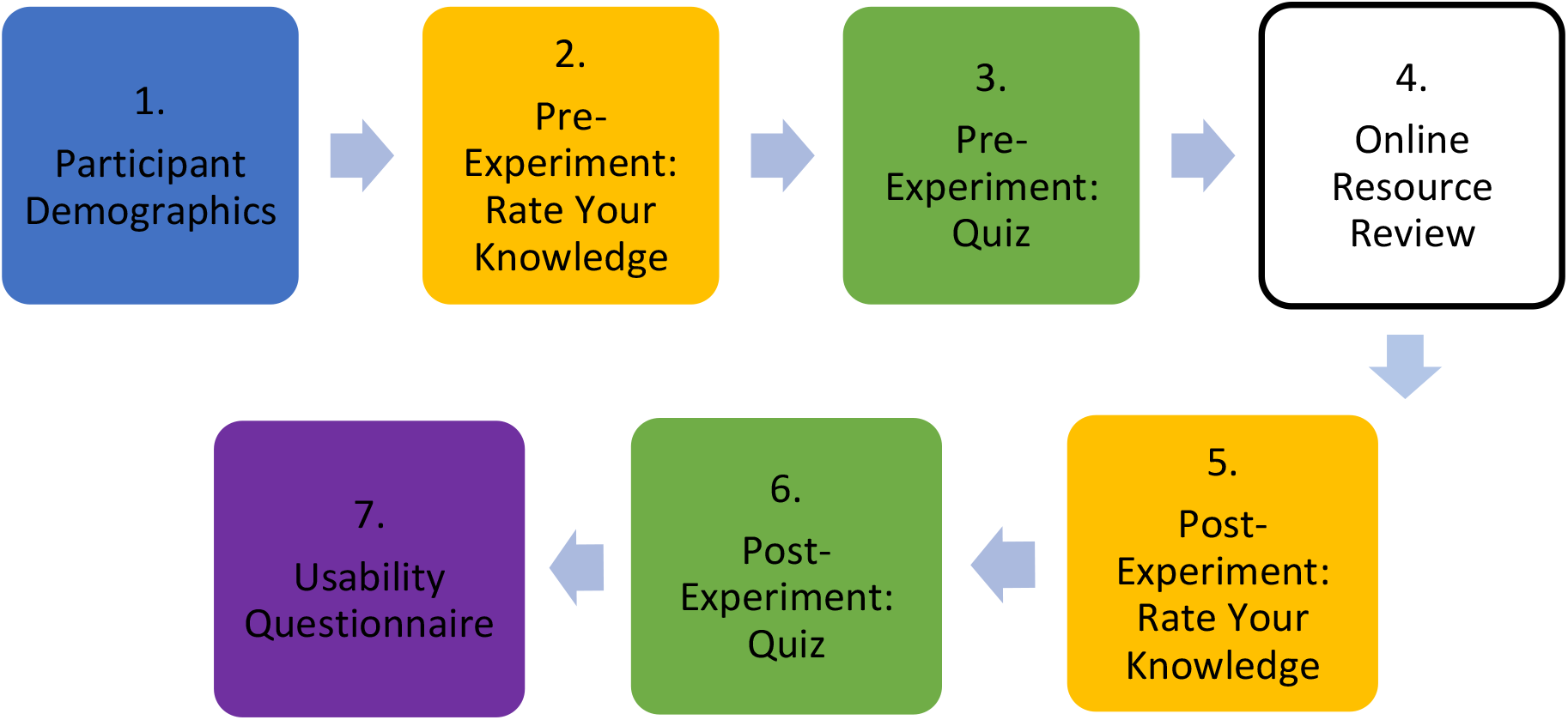
The flow of the seven stages of the experimental questionnaire.

The Online Resource Review stage (Stage 4) differed between the CG and TG, with the CG reviewing the COG-UK/ME and the TG reviewing the SSPME. A 5-point Likert scale was used in Stages 2 and 5 with questions asking the user to rate their level of knowledge on different parts of the SARS-CoV-2 structure, with 1 meaning “strongly disagree” and 5 meaning “strongly agree”; this was used to measure confidence of knowledge. Additionally, the data obtained from the questionnaires (Stages 3 and 6) would allow the researchers to determine the knowledge acquisition of the participants. In order to measure the usability of the resources in this project (Stage 7), the System Usability Scale (SUS) and associated guidelines were used to score the usability of each resource; the SUS score was determined by a standardized series of 10-questions assessing their perceived ability to use the resource successfully (Brooke, 1986; Smyk, 2020). An optional open-ended comment section was provided at the end to obtain additional qualitative feedback which would allow the researchers assess the success of different aspects of the application and identify areas for improvement. Apart from this, the question order in the pre-test and post-test was randomised to remove any potential order effect in the study. The results of these methods were used to assess the overall success of the SSPME compared to the COG-UK/ME, and to draw general conclusions about the use of 3D interactive visualisations in scientific education and communication.

## 3 Results

### 3.1 Use of the SSPME Improves Knowledge Acquisition

We tested for differences in participant knowledge acquisition (determined by analysing the differences in the number of correct responses between pre- and post-experiment). The number of correct responses for the CG and TG between the pre-test and post-test quiz questions is shown in Figure 6. As experimental data were not normally distributed, we used a Mann-Whitney U test to statistically analyse the difference between groups, and a Wilcoxon Signed Rank Test to assess the significance of results within each group.

**Figure 6.**
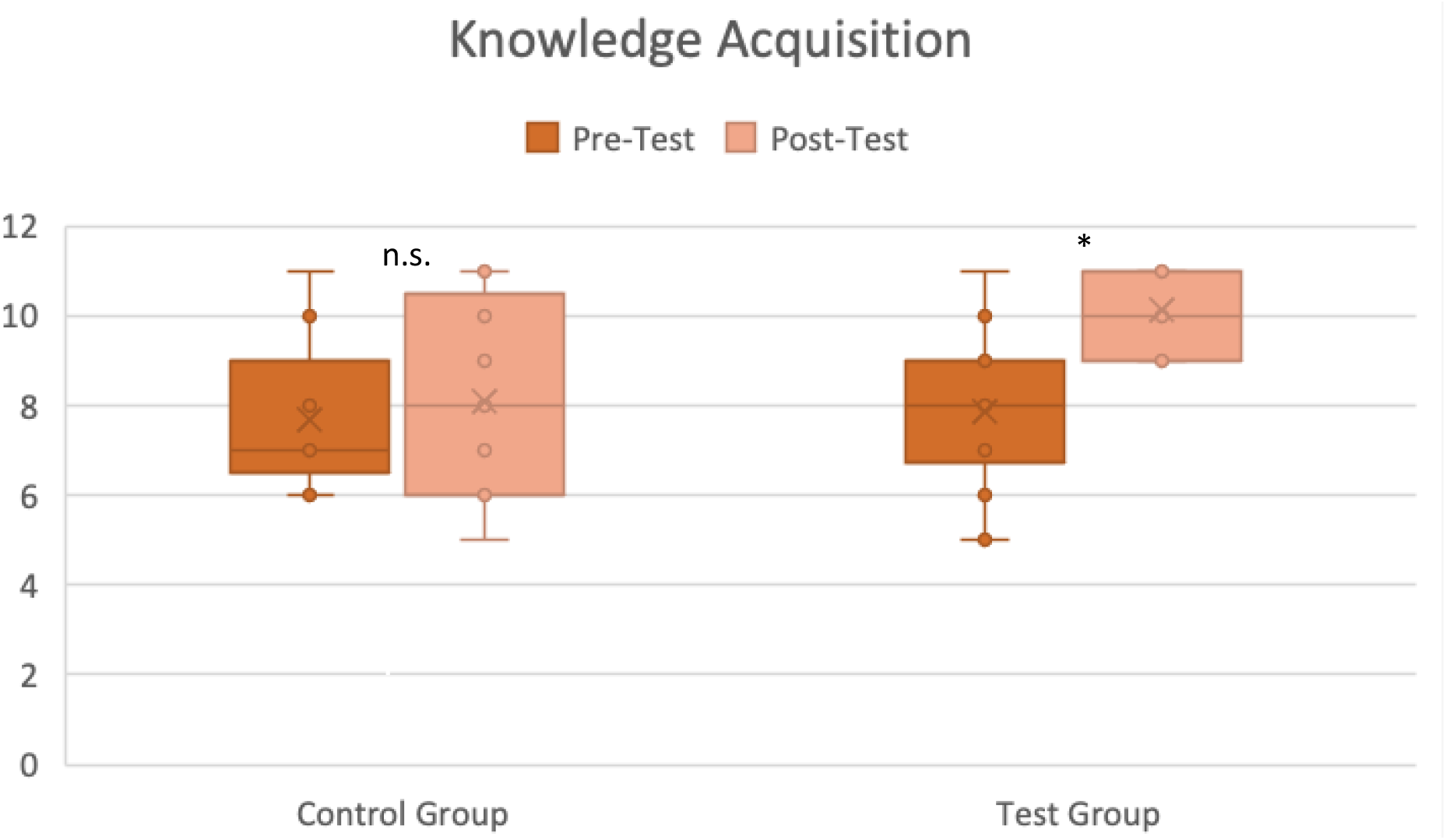
Knowledge acquisition with the CG and TG. The number of correct answers out of 11 questions between the pre-test and post-test quiz questions is shown for both the CG and TG, with individual responses plotted along with box and whisker plots. Differences between pre-test and post-test were tested for significance with a Wilcoxon Signed Rank Test. * p < 0.005; n.s. p > 0.05

We did not find any statistically significant difference between pre-experiment knowledge levels for the CG and TG groups (*U*(*N*_*CG*_ *= 9, N*_*TG*_ *= 14) = 54.00, p = 0.564*). For the CG, there was no statistically significant difference in knowledge acquisition between pre- and post-experiment (*Z*(*N*_*CG*_ *= 9) = -0.79, p = 0.429*). However, for the TG there was a very significant difference in knowledge acquisition between pre- and post-experiment (*Z*(*N*_*TG*_ *= 14) = -3.16, p < 0.005)*. In addition, the post-experiment knowledge acquisition of the TG was significantly higher than that of the CG (*U*(*N*_*CG*_ *= 9, N*_*TG*_ *= 14) = 30.00, z = -2.15, p < 0.05*). These results suggest that CG and TG had similar knowledge before undertaking the experiment, and while the CG did not improve their knowledge after using the COG-UK/ME, the TG acquired knowledge in a significant manner after using the SSPME. Therefore, we concluded that the SSPME was better than the COG-UK/ME in allowing the participants to acquire knowledge after use.

### 3.2 Use of the SSPME Boosts Knowledge Confidence

We tested for differences in participant knowledge confidence (determined by a Likert scale self-evaluation of how well the participant felt they knew the material between pre- and post-experiment) between the CG and TG by Mann-Whitney U test. Before starting the experiment, there were no significant differences in knowledge confidence between the CG and TG. However, following the experiment, the CG and TG knowledge confidences differed significantly on all questions (p < 0.05 for all five questions and p < 0.01 for two of the five questions) indicating an increase in knowledge confidence in the TG. We therefore concluded that the SSPME was better than the COG-UK/ME in allowing the participants to feel more confident in their knowledge after using the SSPME.

### 3.3 The SSPME Demonstrates Strong Usability

SUS scores were calculated after user-testing. The COG-UK/ME received 42.78 (Grade F, or ‘Awful’) and the SSPME received 82.68 (Grade A, or ‘Excellent’), showing a dramatic difference between the resources apparent usability for non-expert audiences (Sauro, 2016; Alathas, 2018). From these results, we concluded that the SSPME was more user-friendly to participants than the COG-UK/ME.

### 3.4 Open-Ended Feedback Indicates that the SSPME Is Positively Received By Users

Open-ended feedback received from the CG was used early in the research timeline in order to inform the development of the SSPME (see Supplementary Material). Much of the feedback from the CG expressed a desire for the COG-UK/ME to contain more basic background information to help in understanding the data on the website. Additionally, it was noted that visual aids would be beneficial for the COG-UK/ME.

Open-ended feedback received from the TG was used to assess the effectiveness of the SSPME, and to identify areas for improvement (see Supplementary Material). The feedback received on the SSPME showed a lot of positive responses, describing the SSPME as “*easy to use*” with “*clear*” and “*understandable*” information and animations. Additionally, the TG provided valuable constructive comments, suggesting potential improvements for future versions of the SSPME, such as redesigning and relocating the animation, and implementing interactive models of specific VOCs.

## 4 Discussion

### 4.1 Research Findings

The COG-UK/ME was developed as a data sharing resource for SARS-CoV-2 genome data, with the stated aim of allowing anyone to follow information over time on important changes in the SARS-CoV-2 genome. Given the technical nature of the data presented, the primary audience of COG-UK/ME was specialists in virology, public health, pharmaceutical industry and bioinformatics. However, the widespread public and media interest in SARS-CoV-2 and its variants during the pandemic meant that it was desirable to make this resource accessible to as wide an audience as possible. By doing so, the implications of SARS-CoV-2 mutations and VOCs on COVID-19 restrictions and public health measures could be better understood by the public, improving the science communication about the pandemic as a whole.

User-testing results show that not only did the SSPME improve the knowledge acquisition of the TG, but it also enabled the TG to be more confident in their knowledge of the material. These results also showed that the COG-UK/ME did not have the same impact on the CG, whose knowledge acquisition and confidence of knowledge did not improve as effectively after using the website. Additionally, results revealed that the SSPME has higher usability, earning a place in the top 10% ranking, while the COG-UK/ME fell in the bottom 10% ranking for usability under the SUS guidelines. This suggests that the SSPME can support a better user experience for non-specialist users, compared to the COG-UK/ME. These results are promising, and indicate that this visual resource could benefit the wider public.

### 4.2 Experimental Review and Limitations

There were limitations placed on the user-testing process of this project due to the COVID-19 pandemic. This project was conducted entirely remotely, which had advantages and disadvantages. It is possible that recruiting participants in an online format via social media allowed a wider participant population to view the study and sign-up. However, it is also possible that conducting the user-testing in person could have increased participation and improved questionnaire follow-through; noting the difference between the number of individuals who initially signed up for the study (N_CG_ = 55, N_TG_ = 61) and the number who fully completed the study (N_CG_ = 9, N_TG_ = 14) suggests that this may be the case. Email reminders were sent to combat this issue during the research, but they did not substantially improve completion. Further work could examine the knowledge retention of participants to determine if the SSPME had lasting impacts on participant knowledge of SARS-CoV-2 spike proteins and VOCs. User-testing could then examine the participants’ knowledge after a longer period of time to assess medium to long-term retention of knowledge. Furthermore, to eliminate any potential order effects on participant knowledge, a crossed-factorial experimental design could be applied during user-testing.

When reviewing the demographics of the participants in this study, we noted that many participants had strong academic backgrounds, with 57% of both the CG and TG participants having a Master’s degree or other higher degree. This level of education and training can be argued as not fully representative of the ‘general public’ that the SSPME was originally designed for. It is likely that by advertising this scientific study primarily through platforms with higher engagement from those interested in science, the participants who signed up for this study already had some background in scientific topics explored in this project. This concern is reflected in Table 1, where many participants considered their knowledge of these topics to be at least “basic”. In the future, conducting participant recruitment across more varied channels and platforms could result in a greater number of participants with more wide-ranging educational backgrounds. However, the importance of advanced research to everyday life during the SARS-CoV-2 pandemic meant that even people with a high level of relevant education and training could benefit from explanatory resources. In this case, even though the TG had well-educated participants, they still demonstrated an increase in both knowledge acquisition and confidence of knowledge after using the SSPME. Therefore, the data obtained in this study is still very encouraging in analysing the potential of the SSPME for the wider public.

### 4.3 Future Research in Visual Scientific Communication

Considering the encouraging user-testing results of this study, future research could explore how implementing interactive 3D visualisations could assist in improving scientific communication on other complex topics. Including this technology into science communication can give the viewer a new perspective, allowing them to visually explore challenging topics such as molecular structures and dynamic processes. Without interactive visualisations, the viewer can only try to grasp this material through text and static images, which can be extremely difficult for understanding topics that require building a complex mental image of something that, due to its microscopic scale, is not part of our normal experience (Jenkinson, 2018; Höffler, 2010; Barak & Hussein-Farraj, 2013). Beyond their educational use, applications of this sort can also have a wider public value. During the COVID-19 pandemic, it has been clear that miscommunication and misinformation can circulate widely with deleterious consequences for public health measures. The development of effective communication tools – like the SSPME – can increase the public’s understanding of complex scientific topics, and by doing so may help them to make well-informed decisions about how to protect themselves and the people around them from viruses such as SARS-CoV-2.

### 4.4 Conclusion

This project successfully developed an interactive 3D visual resource, the SSPME, to allow non-expert audiences to understand SARS-CoV-2 spike mutations and VOCs, demonstrating promising results on the SSPME’s potential for educating the public on these topics. These user-testing results suggest that the COG-UK/ME website will be made more accessible to the general public through the 3D visualisations and interactivity of the SSPME. Not only could this benefit those without a scientific background, but it could also be valuable for informed non-specialist communicators such as teachers and journalists.

## Acknowledgements

Work in the Hutchinson group is funded by the MRC [MR/N008618/1 and MR/V035789/1] and by The University of Glasgow. JH and DLR are funded by the MRC [MC_UU_12014/12]. WTH is funded by the MRC [MR/R024758/1 and MR/W005611/1]. The work described in this article was devised and carried out as part of an MSc in Medical Visualisation and Human Anatomy at The Glasgow School of Art and The University of Glasgow.

## Author contributions

SI: Conceptualisation, Investigation, Visualisation, Methodology, Software, Writing – original draft; WTH: Resources; JH: Resources; DLR: Resources; MP: Supervision, Writing – review and editing, EH: Supervision, Writing – review and editing.

The authors declare no conflict of interest.

## References

Alathas, H., 2018. How to Measure Product Usability with the System Usability Scale (SUS) Score. [Online] Available at: https://uxplanet.org/how-to-measure-product-usability-with-the-system-usability-scale-sus-score-69f3875b858f [Accessed 02 08 2021].

Anon., InPress. Biomedical Visualisation. 12 ed. s.l.:Springer.

Barak, M. & Hussein-Farraj, R., 2013. Integrating Model-Based Learning and Animationsfor Enhancing Students’Understandingof Proteins Structure and Function. Research in Science Education, 43(2), pp. 619–636.

Benton, D. J. et al., 2020. Receptor binding and priming of the spike protein of SARS-CoV-2 for membrane fusion. Nature, 588(7837), pp. 327–330.

Bonner, A. et al., 2009. Location of secretory component on the Fc edge of dimeric IgA1 reveals insight into the role of secretory IgA1 in mucosal immunity. Mucosal Immunology, 2(1), pp. 74–84.

Brooke, J., 1986. SUS: a “quick and dirty” usability scale. In: P. W. Jordan, B. Thomas, B. A. Weerdmeester & A. L. McClelland, eds. Usability Evaluation in Industry. London: Taylor and Francis.

Bruce-Low, S. S. et al., 2013. Interactive mobile learning: a pilot study of a new approach for sport science and medical undergraduate students. Advances in Physiology Education, 37(4), pp. 292–297.

Casalino, L. et al., 2020. Beyond shielding: the roles of glycans in the SARS-CoV-2 spike protein. ACS Central Science, 6(10), pp. 1722–1734.

Chan, J. F.-W. et al., 2020. Genomic characterization of the 2019 novelhuman-pathogenic coronavirus isolated from a patient with atypical pneumonia after visiting Wuhan. Emerging Microbes & Infections, 9(1), pp. 221–236.

Höffler, T. N., 2010. Spatial Ability: Its Influence on Learning with Visualizations—a Meta-Analytic Review. Educational Psychology Review, 22(3), pp. 245–269.

Harvey, W. T. et al., 2021. SARS-CoV-2 variants, spike mutations and immune escape. Nature Reviews Microbiology, Volume 19, pp. 409–424.

Heo, L. & Feig, M., 2020. SARS-Cov-2 Protein Structure Models. [Online] Available at: https://github.com/feiglab/sars-cov-2-proteins/tree/master/Membrane [Accessed 21 May 2021].

Iannucci, S. et al., InPress. Using Molecular Visualisation Techniques to Explain the Molecular Biology of SARS-CoV-2 Spike Protein Mutations to a General Audience. In: L. Shapiro & P. Rea, eds. Biomedical Visualisation (Advances in Experimental Medicine and Biology). s.l.:Springer.

Jenkinson, J., 2018. Molecular Biology Meets the Learning Sciences: Visualizations in Education and Outreach. Journal of Molecular Biology, 430(21), pp. 4013–4027.

Ke, Z. et al., 2020. Structures and distributions of SARS-CoV-2 spike proteins on intact virions. Nature, 588(7838), pp. 498–502.

Letko, M., Marzi, A. & Munster, V., 2020. Functional assessment of cell entry and receptor usage for SARS-CoV-2 and other lineage B betacoronaviruses. Nature Microbiology, 5(4), pp. 562–569.

Peacock, T. P., Penrice-Randal, R., Hiscox, J. A. & Barclay, W. S., 2021. SARS-CoV-2 one year on: evidence for ongoing viral adaptation. Journal of General Virology, 102(4), p. 001584.

Sauro, J., 2016. Measuring Usability With The System Usability Scale (SUS). [Online] Available at: https://www.userfocus.co.uk/articles/measuring-usability-with-the-SUS.html [Accessed 28 07 2021].

Scapin, G. Y. X. P. W. M. M. R. P. J. J. K. R. & Strickland, C., 2015. Structure of full-length human anti-PD1 therapeutic IgG4 antibody pembrolizumab. Nature Structural & Molecular Biology, 22(12), pp. 953–958.

Schmiege, P., Fine, M., Blobel, G. & Li, X., 2017. Human TRPML1 channel structures in open and closed conformations. Nature, 550(7676), pp. 366–370.

Smyk, A., 2020. The System Usability Scale & How It’s Used in UX. [Online] Available at: https://xd.adobe.com/ideas/process/user-testing/sus-system-usability-scale-ux/ [Accessed 20 07 2021].

Terasawa, K. et al., 2016. Lysosome-associated membrane proteins-1 and-2 (LAMP-1 and LAMP-2) assemble via distinct modes. Biochemical and Biophysical Research Communications, 479(3), pp. 489–495.

Woo, H. et al., 2020. Developing a fully glycosylated full-length SARS-CoV-2 spike protein model in a viral membrane. The Journal of Physical Chemistry, 124(33), pp. 7128–7137.

World Health Organization, 2022. WHO Coronavirus (COVID-19) Dashboard. [Online] Available at: https://covid19.who.int/ [Accessed 26 06 2022].

Wright, D. et al., 2021. COVID-19 Genomics UK Consortium (COG-UK) Mutation Explorer. [Online] Available at: http://sars2.cvr.gla.ac.uk/cog-uk/ [Accessed 19 May 2021].

Wrobel, A. et al., 2020. SARS-CoV-2 and bat RaTG13 spike glycoprotein structures inform on virus evolution and furin-cleavage effects. Nature Structural & Molecular Biology, 27(8), pp. 763–767.

Yan, R. et al., 2020. Structural basis for the recognition of SARS-CoV-2 by full-length human ACE2. Science, 367(6485), pp. 1444–1448.

Zhao, J. et al., 2017. Molecular basis for the binding and modulation of V-ATPase by a bacterial effector protein. PLoS Pathogens, 13(6), p. e1006394.

